# “DoC-feeling”: a new behavioural tool to help diagnose the Minimally Conscious State

**DOI:** 10.1101/370775

**Authors:** Bertrand Hermann, Gwen Goudard, Karine Courcoux, Mélanie Valente, Sebastien Labat, Lucienne Despois, Julie Bourmaleau, Louise Richard-Gilis, Frédéric Faugeras, Sophie Demeret, Jacobo D Sitt, Lionel Naccache, Benjamin Rohaut

**Author notes:** These authors contributed equally to this work (Bertrand Hermann, Gwen Goudard, Karine Courcoux). Corresponding author: Dr Benjamin Rohaut, Division of Critical Care & Hospitalist Neurology, Columbia University, 177 Fort Washington avenue, MHB 8 Center, Room 300, New York, NY 10032, USA. **Data sharing statement**: All relevant data (including the raw dataset table) is available. **Contributorship statement (alphabetic order)**: Study concept and design: BR, GG, JB, JDS, KC, LD, LN, LRG, SD and SL. Data collection: BH, GG, KC, LD, MV and SL. Analysis and interpretation of data: BH, BR. Drafting of the manuscript: BH, BR and LN. Critical revision of the manuscript for important intellectual content: BH, BR, JDS, LN and SD. Statistical analysis: BH, BR and LN. Study supervision: BH, BR, GG, KC and MV had full access to all the data in the study and take responsibility for the integrity of the data and the accuracy of the data analysis. BH, GG and KC contributed equally to this work.

## Abstract

**Objectives:** The clinical distinction between vegetative state/unresponsive wakefulness syndrome (UWS) and minimally conscious state (MCS) is a key step to elaborate a prognosis and formulate an appropriate medical plan for any patient suffering from disorders of consciousness (DoC). However, this assessment is often challenging and may require specialized expertise. In this study, we hypothesized that pooling subjective reports of the level of consciousness of a given patient across several nursing staff members can be used to clinically detect MCS.

**Setting and Participants:** Patients referred for consciousness assessment were prospectively screened. MCS (target condition) was defined according to the best Coma Recovery Scale-Revised score (CRS-R) obtained from expert physicians (reference standard). “DoC-feeling” score consisted in the median value of multiple ratings of patient’s behavior observation pooled from multiple staff members during a week of hospitalisation (index test). Individual ratings were collected at the end of each shift using a 100mm visual analog scale, blinded from the reference standard. Diagnostic accuracy was evaluated using area under the receiver operating characteristic curve (AUC), sensitivity and specificity metrics.

**Results:** 692 ratings performed by 83 nursing staff members were collected from 47 patients. Twenty patients were in a UWS and 27 in a MCS. DoC-feeling scores obtained by pooling all individual ratings obtained for a given patient were significantly greater in MCS than in UWS patients (59.2 mm [IQR: 27.3-77.3] vs. 7.2 mm [IQR: 2.4-11.4]; p<0.001) yielding an AUC of 0.92 (95%CI: 0.84-0.99).

**Conclusions:** DoC-feeling capitalizes on the expertise of nursing staff to evaluate patients’ consciousness. Together with the CRS-R as well as with brain imaging, DoC-feeling might improve diagnostic and prognostic accuracy of DoC patients.

**Strengths and limitations of this study:**

- We designed a new behavioural tool called “DoC-feeling” to help face the clinical challenge of the detection of Minimally Conscious State in patients suffering from disorders of consciousness (DoC)
- “DoC-feeling score” quantifies nursing staff’s subjective perception of patient’s consciousness by pooling multiple assessments obtained from multiple caregivers (“wisdom of the crowds”)
- This score which requires no particular training showed a very good accuracy when compared to the gold standard (repeated expert clinical assessment using the Coma Recovery Scale – Revised (CRS-R))
- A validation in a separate cohort would help to determine its place in consciousness assessment
- This score should be tested not only against the CRS-R but also against brain-imaging techniques to test for its capacity to detect covert signs of consciousness

## Introduction

Accurate diagnosis of the level of consciousness in a brain damaged patient is of great importance to better predict recovery. Disorders of Consciousness (DoC) taxonomy has been recently challenged [1–3] but schematically includes the unresponsive wakefulness syndrome (UWS, also termed vegetative state) and the minimally conscious state (MCS). The detection of MCS has a huge prognostic impact since functional outcome is dramatically better for MCS patients [4–8]. However, assessing consciousness in DoC patients can be challenging and in such cases, clinicians may need dedicated clinical tools and brain imaging techniques specifically designed to probe consciousness [9]. Even when using dedicated clinical tools such as the Coma Recovery Scale Revised (CRS-R; [10]), a unique assessment remains associated with a high frequency of diagnostic error [11,12]. To circumvent this limitation, repeated clinical assessments have been proposed but this can be limited by the availability of trained clinicians [13,14].

In this study, we aimed at evaluating the diagnostic accuracy of pooled nursing staff estimations of the level of consciousness in DoC patients. Through their clinical practice, nursing staff (i.e., nurses and nursing assistants) accumulates extended observation time of patient’s behaviour. Interacting with patients through standardized procedures (such as nursing care, medication administration, blood sample, etc…), they spontaneously generate a subjective estimation of the level of consciousness of the patient. Pooling opinions of several individuals have been shown to outperform individual judgements in specific settings (effect known as “wisdom of the crowds”) [15,16]. In this study, we hypothesized that pooling individual nursing staff estimations of the level of consciousness can help in the detection of MCS.

## Methods

### Patients

All patients referred for evaluation of consciousness at the Department of Neurology of La Pitié-Salpêtrière Hospital, Paris, between February 2016 and October 2017 were screened prospectively. On hospital admission patients’ relatives were approached to give consent for participation to the study. All patients with a UWS or MCS condition and consent were eligible.

### Ethics

The protocol conformed the Declaration of Helsinki, the French regulations, and was approved by the local ethic committee (*Comité de Protection des Personnes; CPP n° 2013-A01385-40*; Ile de France 1; Paris, France).

### Evaluation of consciousness

#### Reference standard

Patients were hospitalised in the NeuroIntensive Care Unit and were observed for at least one week during which they encompassed multiple neurological assessments and brain imagery such as high-density EEG, event-related potentials, MRI and PET-scan. Clinical assessments consisted of repeated neurological exams which included the Coma Recovery Scale–Revised (CRS-R; [10]), performed by expert clinicians (BH, BR, FF, LN) belonging to an external expert team in DoC patients. CRS-R scoring ranges from 0 to 23 and is based on the presence or absence of responses on a set of hierarchically ordered items testing auditory, visual, motor, oromotor, communication and arousal function. State of consciousness (i.g., UWS, MCS) is determined by specific key behaviours probed during the CRS-R assessment. For instance, visual pursuit, reproducible movements to command and/or complex motor behaviour scores for MCS[10]. We used the highest level of consciousness among all the CRS-R performed on a given patient as the reference standard. Following this procedure, each patient was thus labelled as being in a UWS or MCS. MCS was the target condition.

Results are expressed in n(%) or median[Inter Quartile Range] as appropriate. ABI: Acute brain injury; CRS-R: Coma Recovery Scale revised; IQR: Inter-Quartile Range; MCS: Minimally Conscious State; NA: Nursing Assistant; Nb: Number; TBI: Traumatic Brain Injury; UWS: Unresponsive Wakefulness Syndrome. * PET-scan was performed only in patients free of mechanical ventilation.

#### Index test

Nursing staff (nurses and nursing assistants) taking care of a DoC patient was asked to fill in a form at the end of their shift containing a scale called “DoC-feeling” (DoC stands for Disorder of Consciousness). DoC-feeling was designed as a 100 mm visual analog scale aiming at quantifying the best patient’s consciousness level observed during the shift. We specifically asked caregivers to rate their “*gut feeling”* about the best level of consciousness observed during the shift or the *“présence” (presence)*, using the French idiom *“le patient est-il là?”* which is very close to the English one *“Is there anybody home?”* (Figure 1; see Figure S1 for the original visual analog scale and its English translation). This wording reproduced the commonly used language to communicate observations relative to consciousness level of a patient among caregivers. Individual DoC-feeling ratings were collected prospectively.

**Figure 1:**
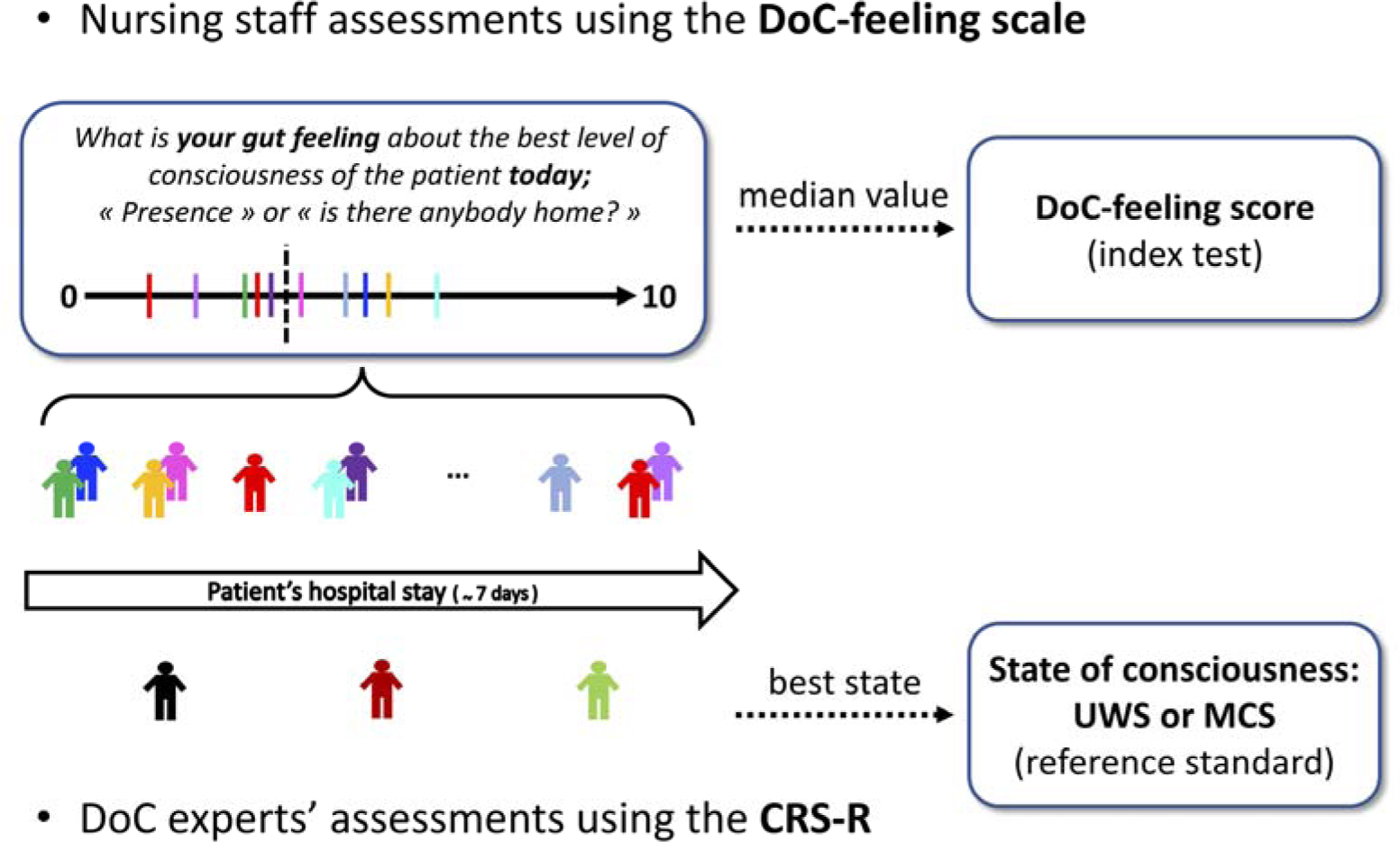
DoC-feeling score. Each patient was evaluated around 3 times by disorders of consciousness (DoC) experts using the Coma Recovery Scale – Revised (CRS-R). In parallel, nursing staff members reported their daily observations using the DoC-feeling visual analog scale. The reference standard was defined as the best state of consciousness observed during one of the CRS-R and the patient was coded as being in a Unresponsive Wakefulness Syndrome (UWS) or a Minimally Conscious State (MCS) accordingly (reference standard). All individual DoC-feeling obtained during the whole hospital stay were pooled and the median value (represented by the vertical dashed line) of the polled results was defined as the DoC-feeling score (index test).

Caregivers were blinded to the previous caregivers’ ratings and to the reference standard (the CRS-R) and expert physicians were blinded to the index test. In order to obtain a final global metric, for each patient, all ratings were pooled using the median to obtain the DoC-feeling score that constituted the index test of this study.

#### Clinical data

Demographics, aetiology and time from the acute brain injury were collected. In addition to CRS-R and DoC-feeling ratings, we also collected complementary metrics (such as the classical distinction between wakefulness and awareness, interaction during nursing and/or painful care) using the same VAS approach (see supplementary material Table S1) as well as the best FOUR-score observed during each shift [17].

#### Statistics

Our primary objective was to evaluate the diagnostic accuracy of the index test called “DoC-feeling score” to detect the target condition (MCS) as defined by the standard reference (best CRS-R).

First, to evaluate the association of individuals’ DoC-feeling ratings with the standard reference, we computed a linear mixed model (LMM) using DoC-feeling individual ratings as the dependent variable, the state of consciousness as the fixed effect explanatory variable and patients as well as raters as random effects. Normality of residuals distribution were assessed by visual inspection. LMM provides the optimal approach in order to take into account the non-independence between DoC-feeling ratings due to the repeated measurements over time at both the patient level (same patient rated by several raters) and the rater level (several ratings by rater).

We next pooled the individual ratings obtained for each patient using the median to obtain the DoC-feeling score (index test). We thus obtained a DoC-feeling score as well as a reference standard label (UWS or MCS) for each patient. We performed a direct comparison of the scores between the two populations using a Wilcoxon-Mann-Whitney test. In order to assess diagnostic accuracy of DoC-feeling scores to detect MCS (target condition), we computed the area under the ROC curve (AUC) and report sensitivities and specificities for several cut-offs of DoC-feeling scores. All statistical tests were two-sided. Categorical variables were expressed as numbers (percentage), quantitative variables as median [interquartile range]. Analyses were performed using the R statistical software version 3.4.1 [18]. LMM was performed using the lme4 package [19]. AUC, sensitivity and specificity with their 95% CI were computed using 2000 stratified bootstrap replicates (AUC) and binomial test (sensitivity and specificity) respectively using the pROC package [20].

The Standards for Reporting Diagnostic Accuracy (STARD) were followed thoroughly [21].

### Patient and Public Involvement

No patients or patients’ relatives were involved in the study design or the management of this study. Results of the study have been released as a preprint on a public repository [22].

## Results

### Patients characteristics

#### Flow chart

Seventy-two patients were eligible during the inclusion period, 23 were not included because of a lack of informed consent from a legal representative. Two patients were excluded because they had been diagnosed as conscious (“Exit-MCS”). Forty-seven patients were included in the analysis (see Figure 2).

**Figure 2:**
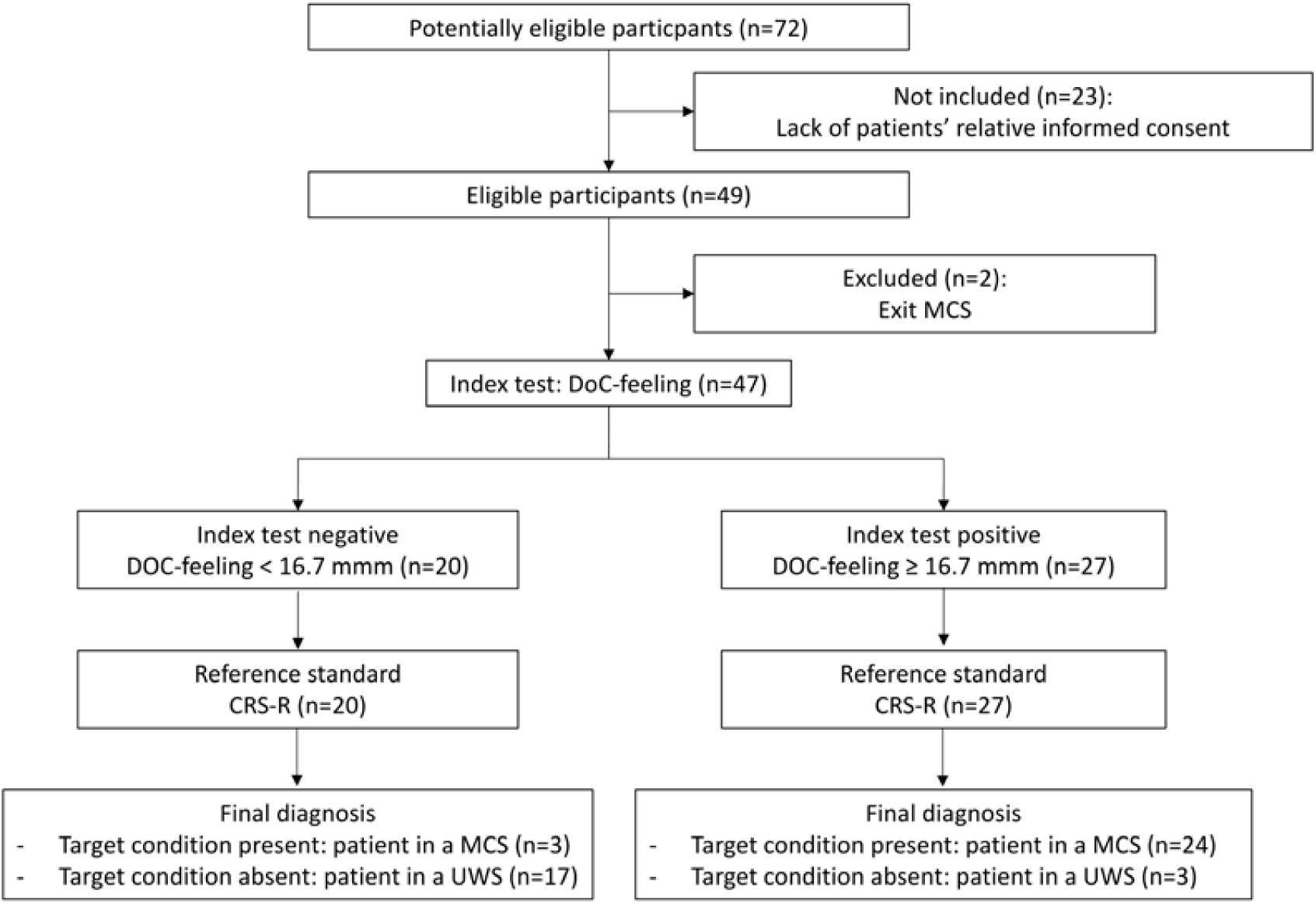
Flow chart. Flow chart representing the repartition of patients while using a DoC-feeling score (index test) cut-off value of 16.7 mm. CRS-R: Coma Recovery Scale – Revised; MCS: Minimally Conscious State; Exit-MCS: Patient able to communicate reliably or to use objects functionally; UWS: Unresponsive Wakefulness Syndrome.

Median age was 49 [32-62] years and 77% (N=36) were women. Main aetiologies of brain injury included anoxia (53%) and traumatic brain injury (17%). Delay between acute brain injury (ABI) and the evaluation was 134 [40-762] days (see Table 1).

**Table 1:**
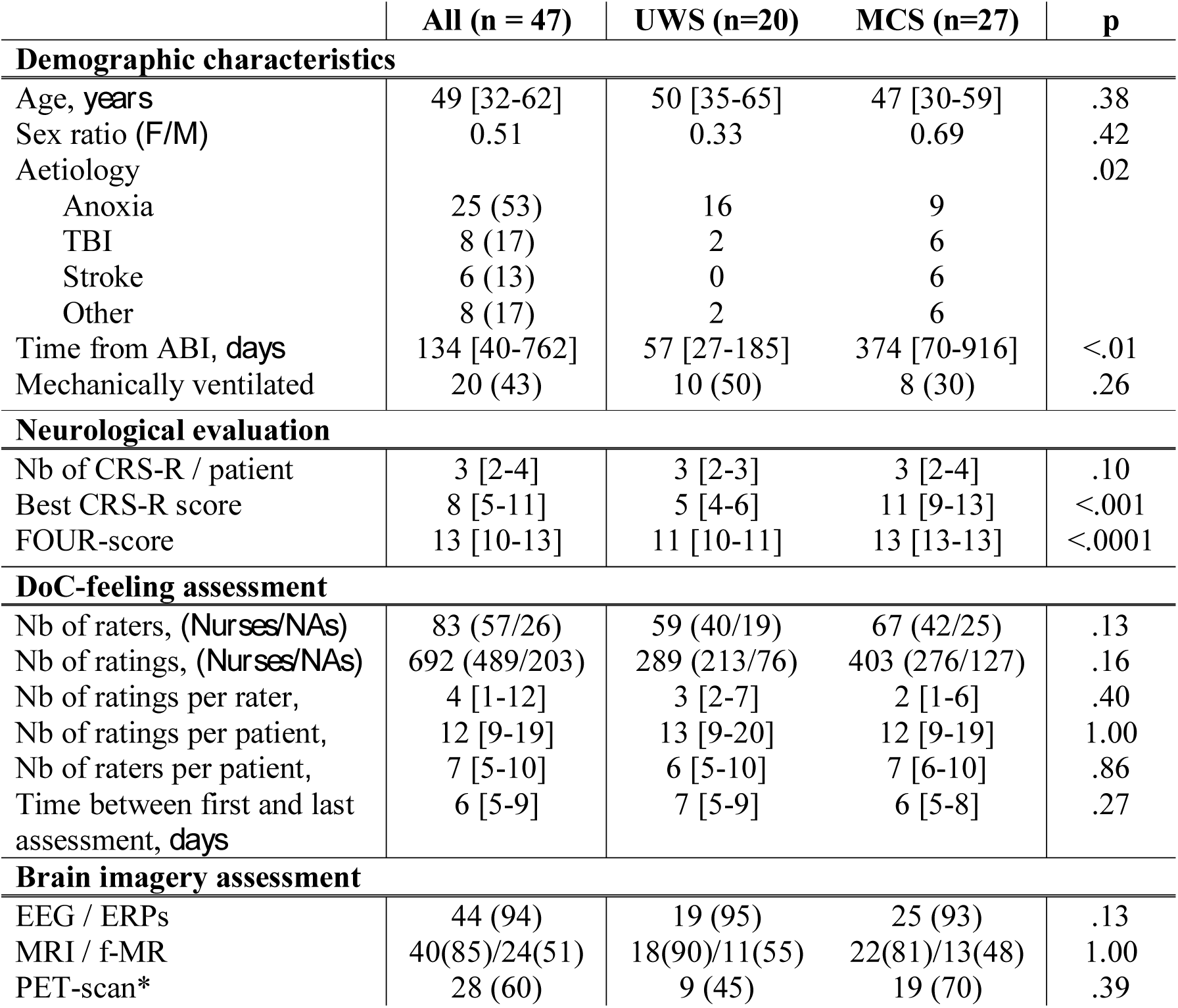
Patient characteristics.

|  | All (n = 47) | UWS (n=20) | MCS (n=27) | p |
| --- | --- | --- | --- | --- |
| **Demographic characteristics** |  |  |  |  |
| Age, *years* | 49 [32-62] | 50 [35-65] | 47 [30-59] | .38 |
| Sex ratio (*F/M*) | 0.51 | 0.33 | 0.69 | .42 |
| Aetiology |  |  |  | .02 |
| Anoxia | 25 (53) | 16 | 9 |  |
| TBI | 8 (17) | 2 | 6 |  |
| Stroke | 6 (13) | 0 | 6 |  |
| Other | 8 (17) | 2 | 6 |  |
| Time from ABI, *days* | 134 [40-762] | 57 [27-185] | 374 [70-916] | <.01 |
| Mechanically ventilated | 20 (43) | 10 (50) | 8 (30) | .26 |
| **Neurological evaluation** |  |  |  |  |
| Nb of CRS-R / patient | 3 [2-4] | 3 [2-3] | 3 [2-4] | .10 |
| Best CRS-R score | 8 [5-11] | 5 [4-6] | 11 [9-13] | <.001 |
| FOUR-score | 13 [10-13] | 11 [10-11] | 13 [13-13] | <.0001 |
| **DoC-feeling assessment** |  |  |  |  |
| Nb of raters, (*Nurses/NAs*) | 83 (57/26) | 59 (40/19) | 67 (42/25) | .13 |
| Nb of ratings, (*Nurses/NAs*) | 692 (489/203) | 289 (213/76) | 403 (276/127) | .16 |
| Nb of ratings per rater, | 4 [1-12] | 3 [2-7] | 2 [1-6] | .40 |
| Nb of ratings per patient, | 12 [9-19] | 13 [9-20] | 12 [9-19] | 1.00 |
| Nb of raters per patient, | 7 [5-10] | 6 [5-10] | 7 [6-10] | .86 |
| Time between first and last assessment, *days* | 6 [5-9] | 7 [5-9] | 6 [5-8] | .27 |
| **Brain imagery assessment** |  |  |  |  |
| EEG / ERPs | 44 (94) | 19 (95) | 25 (93) | .13 |
| MRI / f-MR | 40(85)/24(51) | 18(90)/11(55) | 22(81)/13(48) | 1.00 |
| PET-scan* | 28 (60) | 9 (45) | 19 (70) | .39 |

#### Reference test

One-hundred and forty-seven CRS-R assessments were performed, with a median of 3 [2-4] per patient (ranging from 2 to 6). According to the best CRS-R (reference standard), 27 patients (57%) were diagnosed as being in a MCS (target condition) and 20 (43%) were classified as being in a UWS. Patients with MCS less frequently had suffered from anoxia and had a longer delay between the acute brain injury and the study assessment (see Table 1). No differences were found in the number of CRS-R assessment per patient or brain imaging explorations between UWS and MCS patients.

#### Index test

Six-hundred and ninety-two DoC-feeling individual ratings were obtained (median of 12 [9-19] ratings per patient). 83 caregivers, 57 Nurses and 26 NAs (47 neuro-ICU staff members and 36 float staff members) participated to the study. Each nursing staff member filled a median of 4 [1-12] evaluations. Median delay between the first and the last individual rating was 6 days [5-9]. No statistical differences were found between UWS and MCS in terms of number of DoC-feeling ratings per patient, number of rater per patient or in terms of number of ratings per rater.

### Analysis of individuals DoC-feeling ratings

Inspection of the 692 DoC-feeling ratings’ distribution revealed higher values for MCS than for UWS patients but with an important variability of ratings for a given patient (Figure 3). The linear mixed model (LMM) analysis revealed a strong significant association between DoC-feeling individual ratings and the state of consciousness (t-value=6.47, df=45, p<0.001).

**Figure 3:**
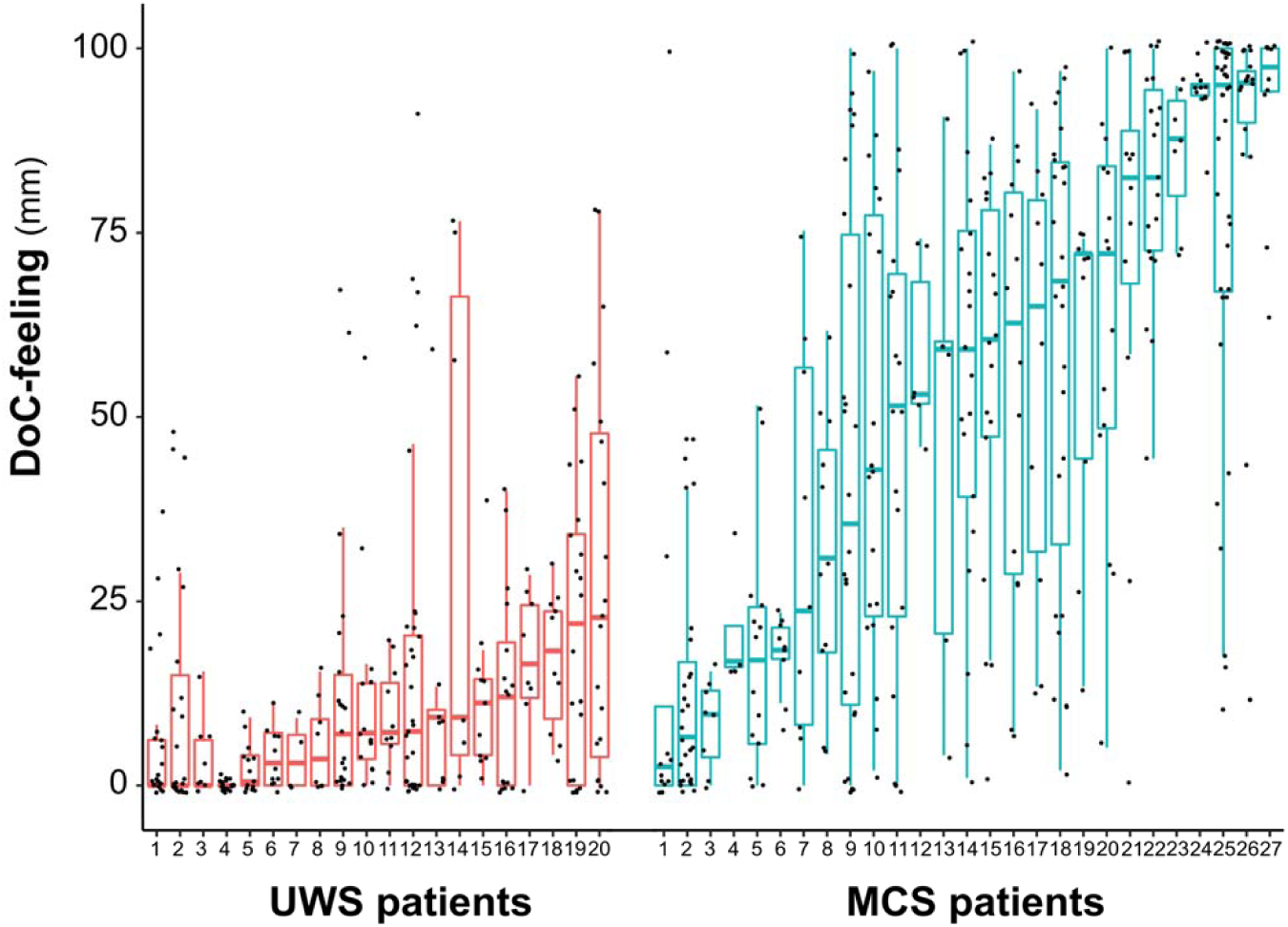
Individual DoC-feeling ratings. DoC-feeling ratings tended to be smaller in UWS patients when compared to MCS patients. All measures are presented (n=692) alongside boxplots helping to visualize the median and the interquartile ranges for both UWS (on the left in red) and MCS (on the right in turquoise) patients.

### Diagnostic accuracy of DoC-feeling scores

Overall, patients underwent 12 [9-19] DoC-feeling individual ratings performed by 7 [5-10] different raters. All DoC-feeling ratings obtained for a given patient were summarized using the median to obtain a pooled metric called DoC-feeling score (index test, Figure 4 a). Median DoC-feeling scores were smaller for UWS than for MCS patients (7.2 mm [2.4-11.4] vs. 59.2 mm [27.3-77.3] respectively; p<0.001; Figure 4b). ROC curve revealed excellent accuracy at detecting MCS (AUC=0.92 [95% CI 0.84-0.99]; Figure 4c) with, for instance, a sensitivity of 89% [95% CI 71-98] and a specificity of 85% [95% CI 62-97] when using a DoC-feeling score cut-off at 16.7 mm (Figure 4d). Note that this cut-off is only used to give the reader an idea about the diagnostic performances using the more intuitive sensitivity and specificity metrics (see discussion). The 6 misclassified patients using this cut-off are presented in the supplementary material (supplementary results and Table S3). Simulations of AUCs using an increasing number of rating per patient suggest that a number of 4 ratings is needed to reach an 0.9 AUC (supplementary material Figure S3).

**Figure 4:**
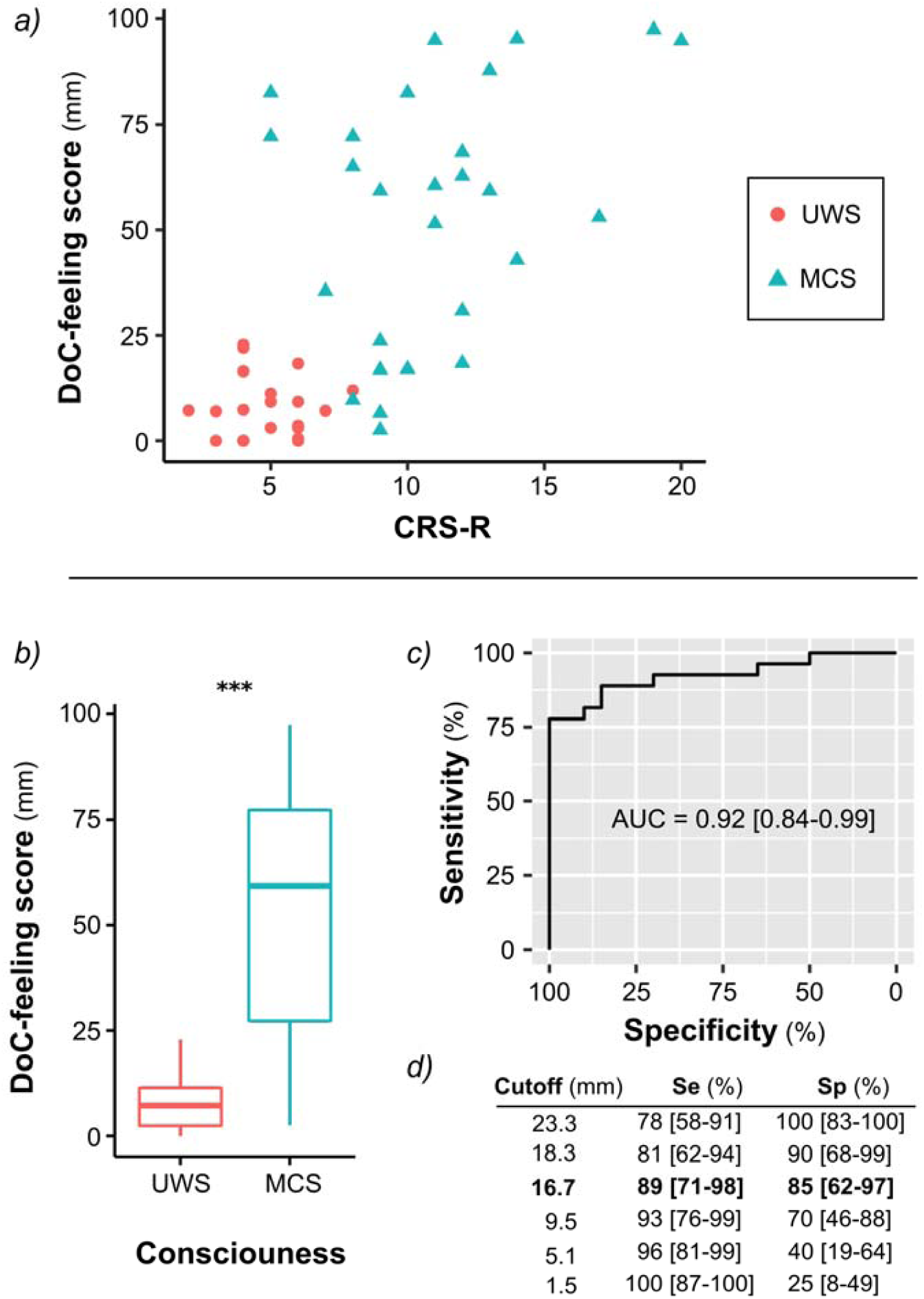
DoC-feeling scores. Median of the individual ratings obtained for each patient were pooled to obtain one DoC-feeling score per patient. DoC-feeling scores tended to be smaller for UWS than for MCS patients (a, b) and to correlate with the total of the CRS (a). Area under the ROC curve (c), sensitivity (Se) and specificity (Sp) for several cutoffs (d) revealed very good performances at identifying the MCS. ***: p<0.001.

## Discussion

In the present study we developed and assessed a new behavioural tool called DoC-feeling to help diagnose MCS. This score which pools repeated measures obtained among several caregivers over several days of evaluation showed a very good accuracy to diagnose MCS.

While this tool is not intended to replace the clinical examination nor the current CRS-R gold standard, we propose it as a complementary one. Indeed, we believe that, taking advantage of valuable information collected by all caregivers involved in the care of a DoC patient, the implementation of DoC-feeling could improve the overall diagnostic accuracy of DoC patients. Caregivers are trained to evaluate pain and suffering in patients during all delivered procedures. These procedures constitute standardized interactions that can allow the generation of very reliable heuristic processes to assess one’s percept in term of pain suffering and also consciousness.

Pooling opinions of several individuals has been previously shown to outperform individual judgements in specific settings. Recently, there has been a growing interest for this kind of approach (called collective intelligence or “wisdom of the crowds”) in the medical field, especially in diagnosis procedure (diagnosis of skin cancer, mammography screening, etc …) [23,24,15,25].

In that perspective, quantifying expertise that is not restricted to physicians might be of prime interest. Capitalizing on assessments of consciousness gathered at any hour of the day and through multiple observers may also potentially increase our ability to detect signs of consciousness in these patients who usually show large fluctuations of cognitive state and of arousal[13]. This tool may also help to better describe and quantify these fluctuations. Additionally, this tool also enables to acknowledge the caregiver group expertise and to increase care team attention through a coherent and cumulative set of observational data.

The good accuracy of DoC-feeling obtained in our setting is likely to be generalizable elsewhere. First, as the distribution of CRS-R scores obtained in this cohort spanned most of the possible CRS-R scores, it is unlikely that the good accuracy of DoC-feeling can be the result of two easily discernible patients’ clusters. Second, as all the patients included in this study, either in an acute or a chronic stage, were specifically referred to our institution for expertise, it is most likely that our cohort was actually representative of patients for whom the diagnosis is the most difficult. However, we would like to emphasise that the optimal cut-off might be variable across teams, according to the overall group bias.

Our study presents some limitations inherent to the aim of developing a pragmatic and easily implementable tool in daily clinical practice. First, as for all study on consciousness disorders, we faced a typical situation of imperfect gold standard. Although CRS-R is still the more widely accepted reference, the optimal number of assessments remain unknow[14]. According to a recent study, using 3 CRS-R assessments can lead to a 17% rate of misdiagnoses [13]. It is worth noting that this is exactly the reason why we developed DoC-feeling. CRS-R requires a specialized expertise that is not available everywhere and that can be extremely time-consuming, especially now that multiple assessments are recommended[14]. In sharp contrast, DoC-feeling scale could be implemented in any team, is much faster and allows multiple observations per day easily. Second, in addition to their clinical observations, caregivers might have been influenced by other factors that would have been very difficult to control. For instance, caregivers might have been influenced by insights from other caregivers or, in case of multiple ratings for a given rater, by their previous ratings. However, the variability of individual ratings for a given patient (that tended to increase over time, see supplementary material results) suggests that caregivers did report their own perception independently from each other and their eventual previous ratings. Moreover, interaction in small groups could have actually had a positive effect since pools of small groups insights’ have been shown to outperform the overall judgment of the group [26]. This kind of tool might be less prone to individual subjective bias that can be observed during decision making under high degree of uncertainty such as assessment of DoC patients [27]. Caregivers could have also been biased by the perception of patients’ relatives, although it is commonly acknowledged that relatives frequently lack objectivity (in both directions) in such dramatic situations [12]. Caregivers’ judgments could have been biased by classical predictors of consciousness recovery such as aetiology or delay from ABI. Finally, although the number of involved float staff members and the result of a preliminary survey assessing prior knowledge of regular nursing staff on DoC (supplementary material Figure S2 and Table S2) suggest together that DoC-feeling should be accurate in other settings, the monocentric design of this study requests external validation.

Despite these limitations, we think that the implementation of DoC-feeling score can significantly improve diagnostic accuracy and confidence in the diagnosis when supporting other metrics (i.e., CRS-R and functional brain imaging at rest or during cognitive tasks). Moreover, when incongruent with other metrics, DoC-feeling score could also be useful. Indeed, this could either suggest that clinical elements have been missed by physicians while performing punctual CRS-R assessments but it could also reveal, in case of discrepancy with all the other markers (clinical and brain imagery), a possible misperception of patient’s consciousness level that need to be acknowledged and considered in any further medical decision processes. This last point could be crucial in bridging the gap between the caregiver’s team and the patient’s relatives in situations of conflict.

In conclusion we propose a new behavioural tool called DoC-feeling that can help in improving the diagnostic accuracy of MCS and thus promote better prognostic and decision making for DoC patients.

## Acknowledgments

We thank all the members of the Pitié-Salpétrière hospital Neuro-ICU led by Sophie Demeret (Medical Director), Julie Bourmaleau and Louise Richard-Gilis, (Head Nurses) who have participated in this study (alphabetic order): Jérémie Abitbol, Fatiha Ait Yata Azzi, Binta Bah Fatoumata, Francis Bolgert, Sandrine Briand, Sandra Coelho, Alexia Camuzat, Marie-Chantal Colmar, Flora Cherruault, Cecile Chordi, Véronique Cottin, Bintou Coulibaly, Karine Courcoux, Mélanie Dalibard, Lucienne Despois, Estelle Dumarey, Atef El Ouarghi, Helene Espiand, Cécilia Eltebert, Fabrice Fanhan, Agnès Flament, Suzelle Fontano-Marie, Pascale Fournier, Céline Frammezelle, Gwen Goudard, Alexandra Grinéa, Nouara Harchaoui, Marie Harmancij, Claire Jacqueminet, Charlotte Janvier, Jamila Kebli, Bouchra Khedaoui, Sébastien Labat, Aurélie Lemoal, Kim Louis-Joseph, Brice Lucas, Valérie Maes, Sophie Maillard, Romain Maurel, Madely Petit, Floriane Pépin, Isabelle Picot, Eva Proneur, Manuela Roselmac, Sylviane Saintini, Mélody Seidel, Yolène Sully, Kelly Tcha, Laura Verbaux, Nicolas Weiss, Kelly Yanganju.

We thank all the members of the PICNIC-Lab “DoC-Team”, dedicated to the improvement of care of patients suffering from disorder of consciousness led by Lionel Naccache (alphabetic order): Athena Demertzi; Denis Engemann; Frédéric Faugeras; Bertrand Hermann; Pauline Pérez; Federico Raimondo; Benjamin Rohaut; Johan Stender; Mélanie Valente and Jacobo Sitt.

We thank Raphael Porcher for his help on statistical issues and Jan Claassen for his final review of our manuscript.

